# Human Cerebral Spheroids Undergo Activity Dependent Changes In Cellular Composition And Microrna Expression

**DOI:** 10.1101/2022.02.04.478858

**Authors:** Thomas Parmentier, Fiona James, Elizabeth Hewitson, Craig Bailey, Nicholas Werry, Steve D. Sheridan, Roy H. Perlis, Melissa Perreault, Luis Gaitero, Jasmin Lalonde, Jonathan LaMarre

## Abstract

Activity-induced neurogenesis has been extensively studied in rodents but the lack of ante mortem accessibility to human brain at the cellular and molecular levels limits studies of the process in humans. Using cerebral spheroids derived from human induced pluripotent stem cells (iPSCs), we investigated the effects of increased neuronal activity on neurogenesis. Our studies demonstrate that increasing neuronal activity with 4-aminopyridine in 3-month-old cerebral spheroids is associated with increases in the numbers of new neurons and decreases in the population of new glial cells. We also observed a significant decrease in the expression of miR-135a, which has previously been shown to be decreased in exercise-induced neurogenesis. Predicted targets of miR-135a include key participants in the SMAD2/3 and BDNF pathways. Together, our results suggest that iPSC-derived cerebral spheroids are an attractive model to study some aspects of activity-induced neurogenesis.

## INTRODUCTION

Human brain development is a highly orchestrated process during which precursors of neuronal and glial cells organize, as they differentiate, into specialized cellular circuits required for normal function. Because neurogenesis and gliogenesis in humans occur primarily *in utero*, studying these processes, and understanding how they may contribute to the onset and progression of certain neurological disorders, is particularly challenging. The discovery that neurogenesis continues in certain discreet regions of the adult brain, in particular the dentate gyrus of the hippocampus, has improved our knowledge of the factors that modulate neurogenesis and the functional implications of adding new neurons to existing neuronal networks (Deng et al., 2010; Moreno-Jiménez et al., 2019; Tashiro et al., 2007). Interestingly, postnatal hippocampal neurogenesis is enhanced by different stimuli that increase activity, such as hippocampal-dependent learning tasks, long-term potentiation, exposure to an enriched environment, and motor activity (Bruel-Jungerman et al., 2006; Gould et al., 1999; van Praag, 2005). Epileptic seizures, which are uncontrolled events where the brain undergoes a pathological increase in neuronal activity, have also been found to increase the formation of new neurons (Jessberger et al., 2007). This activity-dependent postnatal neurogenesis phenomenon is attributed to both an increase in proliferation and differentiation of neural stem cells and an increase in the survival of newly formed neurons (Bruel-Jungerman et al., 2006; Tashiro et al., 2007).

The molecular underpinnings of activity-dependent neurogenesis, and how they relate to human brain development *in utero*, are not well understood. Neural stem cells have been shown to respond to neurotransmitters and growth factors released by neurons which then can influence the proliferation and differentiation of these neural stem cells both *in vitro* and *in vivo* (Bao et al., 2017; Botterill et al., 2015; Deisseroth et al., 2004; Tozuka et al., 2005). Because of inherent limits in probing the developing human brain, and the scarcity of human fetal brain samples, the cellular and molecular mechanisms of activity-dependent neurogenesis as well as the scope of this process in human brain development are not well known. Recently, cerebral organoids, a new model of human brain development has been described using human pluripotent stem cells and was shown to recapitulate key milestones of human brain development (Eiraku et al., 2008; Gordon et al., 2021; Lancaster and Knoblich, 2014; Paşca et al., 2015). Cerebral organoids contain most of the diversity of cell types that are present in the developing human brain and self-organize similarly in this model (Paşca et al., 2015; Quadrato et al., 2017; Tanaka et al., 2020; Velasco et al., 2019). Moreover, with increasing time in culture, cerebral organoids generate complex oscillatory activity that cannot be recapitulated with more simple culture methods such as neurospheres that are 3D aggregates lacking the layered organization of cerebral organoids (Trujillo et al., 2019). Given their ability to capture many key features of human brain development, cerebral organoids have recently been used to model several neurodevelopmental disorders *in vitro* and study the effect of impaired neurogenesis on neuronal network activity (Birey et al., 2017; Sun et al., 2019; Trujillo et al., 2021). However, it is unknown whether activity-induced neurogenesis can be recapitulated in human cerebral organoids. In this study, we explore the effects of 4-aminopyridine (4AP) — an organic compound known to increase neuronal activity in *ex vivo* brain slices — on the neuronal activity in cerebral organoid and its global effect on neurogenesis. In addition, we examine the changes in specific microRNAs (miRNAs) that are associated with neurogenesis *in vivo*. Overall, our study provides a platform for the investigation of activity-dependent neurogenesis in a complex model of human brain development.

## RESULTS

### Recapitulation of neurogenesis and gliogenesis in cerebral spheroids

Different protocols have been used to generate cerebral organoids from human pluripotent stem cells. To help ensure consistency in cerebral organoid generation in between cell lines and batches, we used a guided protocol that was designed to result in neurons demonstrating a cortical phenotype. This protocol has been shown to reliably generate cerebral organoids termed “cortical spheroids” by dual SMAD inhibition to direct pluripotent stem cells to a neuroectodermal phenotype and then neurons with a dorsal forebrain identity (Yoon et al., 2019). We first verified the development of spheroids by employing immunofluorescence identification of cell types using three different iPSC lines derived from neurologically normal individuals. Embryoid bodies were successfully generated from all 3 iPSC lines and the resulting spheroids progressively increased in size during culture (Supplemental Figure 1). Upon dual SMAD inhibition, cells expressed the neuroectodermal cell marker SOX2 and organized into polarized neural tube-like structures (Figure 1A). Neural progenitor cells expressing the marker Nestin and proliferative marker Ki67 were present in these neural tube-like structures and were radially organized (Figure 1A-C). Neural progenitor cells progressively differentiated into neuroblasts expressing the marker DCX as well as post-mitotic neurons expressing MAP2 and NeuN localized at the periphery of the neural tube-like structures. Glial cells expressing GFAP appeared around day 60 and progressively increased in number (Figure 1A and 1D).

**Figure 1:**
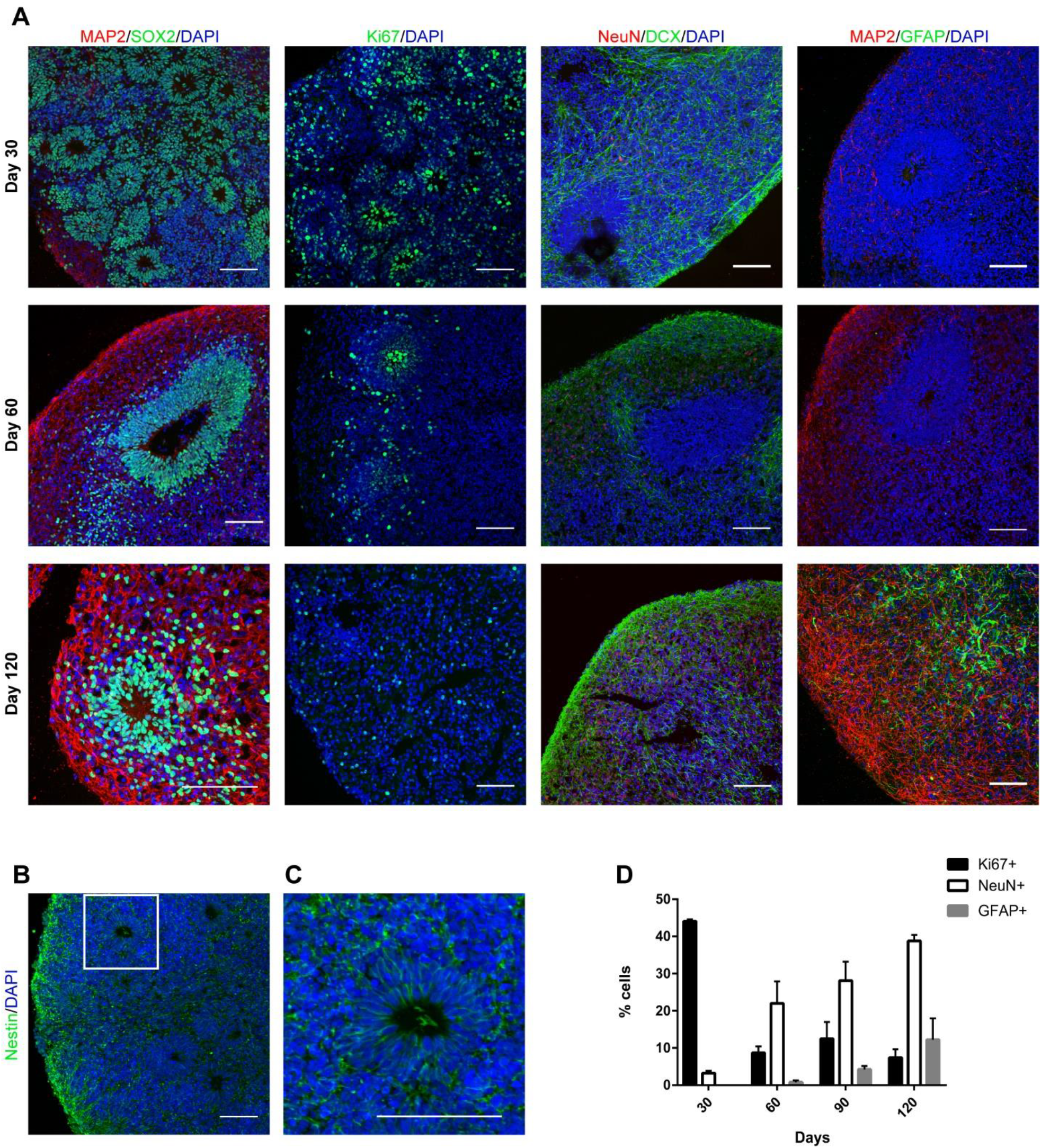
Neurogenesis and gliogenesis in cerebral spheroids. (A) Representative immunofluorescence images of cerebral spheroids after 30, 60 or 120 days of culture. Cells were stained for neuroectodermal cell (SOX2), proliferative cell (Ki67), NSCs (Nestin), immature neuron (DCX), post-mitotic neuron (MAP2, NeuN) or glial cell (GFAP) markers. Bars = 100 μm (B) NSCs express the neural stem cell marker Nestin. A spheroid at day 30. Bar = 100 μm (C) Enlargement of white square in (B) showing that Nestin-positive NSCs are radially organized in a neuroepithelium-like structure. Bar = 100 μm (D) Quantification of cells expressing specific markers for proliferation (Ki67), neurogenesis (NeuN) and gliogenesis (GFAP). Data represent mean +/− SEM of 2-3 organoids/cell line, 3 different iPSC lines.

To verify that putative neurons within the organoids also acquired relevant functional properties, we performed whole-cell recordings of putative neurons in cerebral spheroids between maturation days 70-80 and days 115-120. Functional neuronal maturation was observed through an increase in action potential amplitude, a decrease in membrane resistance and a trend towards an increase in rheobase as neurons matured (Table 1). Furthermore, all excitable cells that were subjected to recording between days 70-80 developed a depolarization block when injected 100 pA (3/3) whereas only 2/8 excitable cells developed a depolarization block after the same stimulation at day 115-120 (Figure 2). Unfortunately, the whole-cell recording procedure proved particularly challenging under these circumstances with 3/11 and 8/21 cells that were successfully acquired were excitable at days 70-80 and 115-120 respectively (Table 1). This relatively low success rate may be attributable to low cell survival after the slicing procedure, immaturity of the neurons in spheroids at these time points and the technical difficulty of whole-cell electrophysiology. Nevertheless, several neurons with mature electrophysiological properties were consistently acquired and recorded, confirming the reliability of the chosen protocol.

**Table 1:**
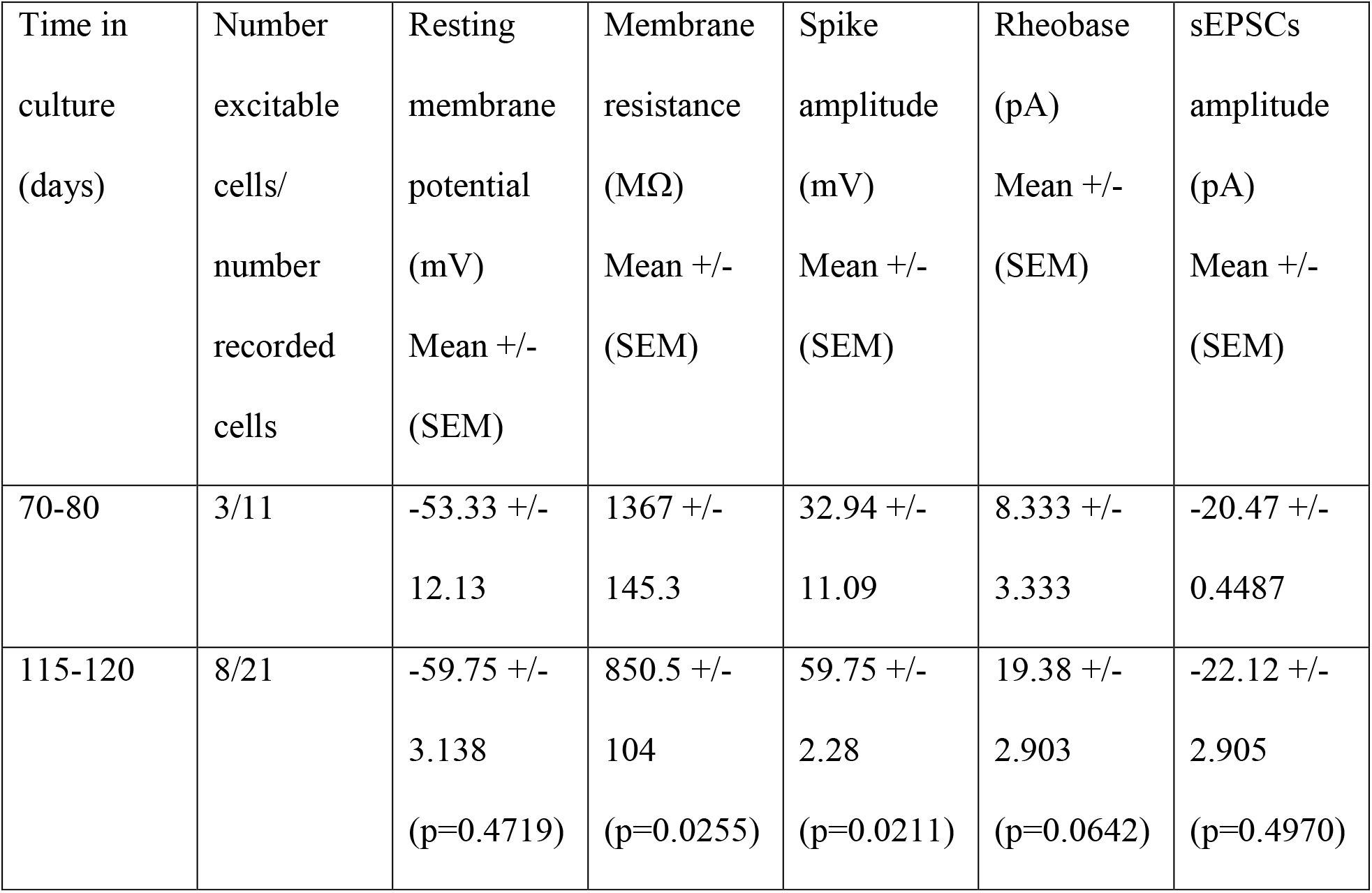
Electrophysiological properties of patched cells across time in culture. Summary of electrophysiological characteristics of excitable cells in cerebral spheroids recorded with whole-cell recordings. Student’s *t*-test.

**Figure 2:**
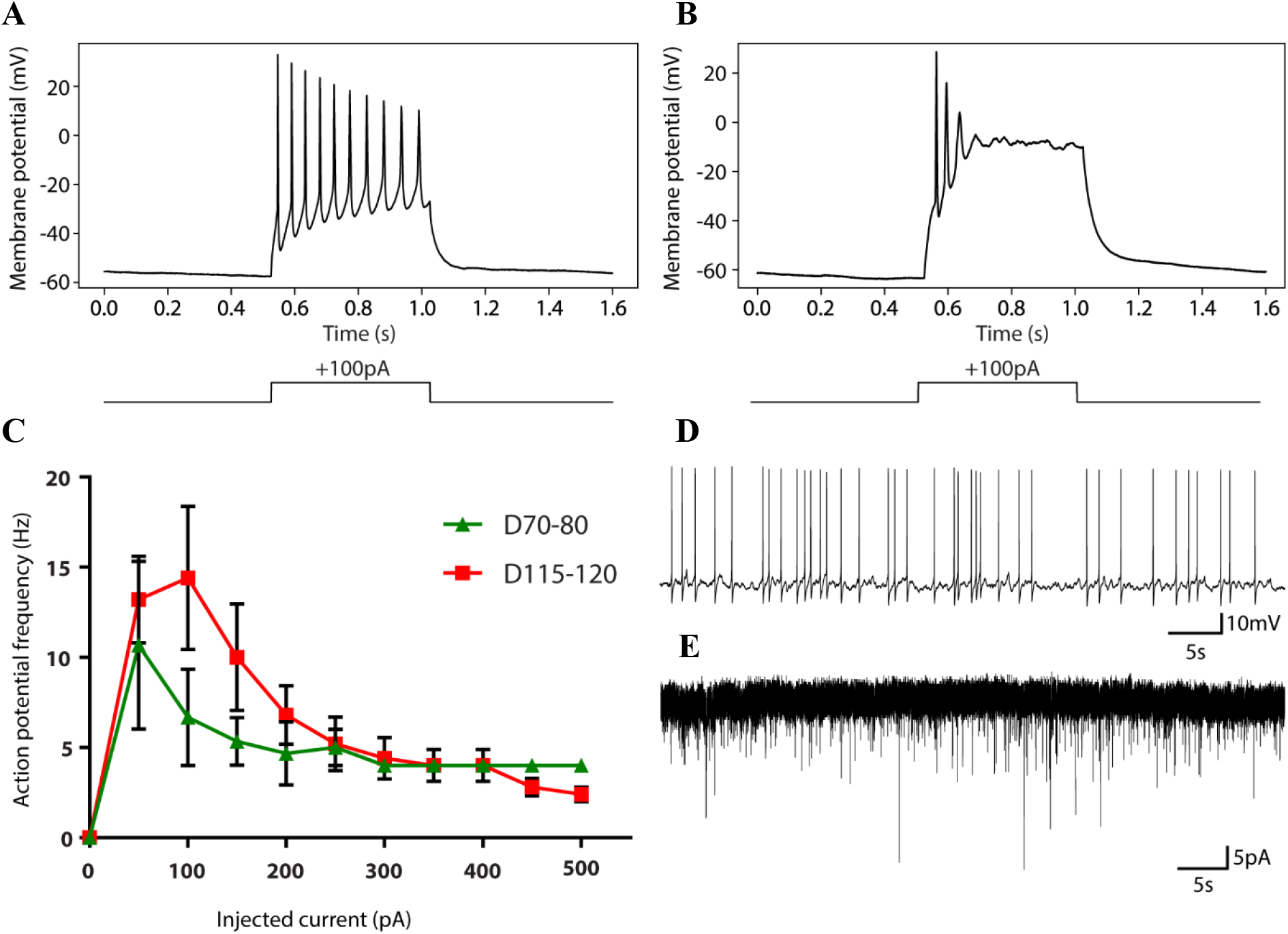
Electrophysiological properties of neurons in cerebral spheroids. (A) Example of a burst of action potentials in a putative mature neuron triggered by100 pA of injected current. (B) Example of a depolarization block in a putative immature neuron. (C) Input-output curves as a measure of excitability in spheroids at different times of culture. Neurons in spheroids show a maximum action potential frequency around 50-100 pA of injected current after which a depolarization block is observed. Data represents mean +/− SEM of excitable cells described in Table 1. (D) Representative trace in current clamp mode showing spontaneous action potentials. (E) Representative trace in voltage-clamp mode showing spontaneous putative sEPSCs.

### Effect of 4AP on neuronal excitability in cerebral spheroids

To investigate whether we could increase neuronal activity in 90-day old cerebral spheroids, we incubated them with 4AP which has been shown to increase neuronal activity in brain slices and to induce epileptic seizures *in vivo* (Gonzalez-Sulser et al., 2011; Heuzeroth et al., 2019; Hsiao et al., 2015). We first followed neuronal activity in individual cells with fluorescent calcium imaging. As expected, 4AP treatment resulted in an approximately 4-fold increase in the number of active cells (67.1956% vs 17.2247%) and in the frequency of calcium waves (71.7704 mHz vs 16.1405 mHz) in active cells of spheroids (Figure 3A-C). Calcium waves had larger mean amplitudes but shorter mean durations in cells of 4AP-treated spheroids compared to controls (Figure 3D-E). To assess neuronal activity with higher temporal resolution and evaluate synchronization within spheroids, we placed individual 90-day old spheroids directly on multi-electrode arrays (MEA). MEA recordings also showed increases in the mean spike firing rate after exposure to 4AP, although it did not reach statistical significance, likely due to variability in the firing rate between individual spheroids (Figure 3F-G). In addition, spheroids demonstrated bursts of activity after exposure to 4AP, which were not observed prior to exposure (electrode burst average frequency 19.3 mHz). These bursts had an average duration of 199ms +/− 2 ms (Figure 3H). However, neuronal activity remained unsynchronized even after stimulation with 4AP (mean synchrony index: 0.0040 and 0.0126 before and after 4AP exposure respectively). Overall, 4AP increased neuronal activity in 90-day-old cerebral spheroids. We verified that cerebral spheroids could form mature neuronal networks with synchronization by repeating the MEA recordings on 160-day-old spheroids (Figure 4). Compared to 90-day-old spheroids, 4AP increased synchronization of activity in 160-day-old spheroids by approximately 12-fold without reaching statistical significance (synchrony index: 0.0062 vs 0.0755 before and after 4AP exposure respectively). Interestingly, while network bursts were not observed in unstimulated 160-day-old spheroids, exposure to 4AP triggered the appearance of synchronized network bursts in 3/4 spheroids with a frequency of 109.9 +/− 4.9 mHz (Figure 4). These network bursts had a duration of 1.5 +/− 0.4 s and during the bursts, spike frequency was 611.9 Hz +/− 217.8 Hz.

**Figure 3:**
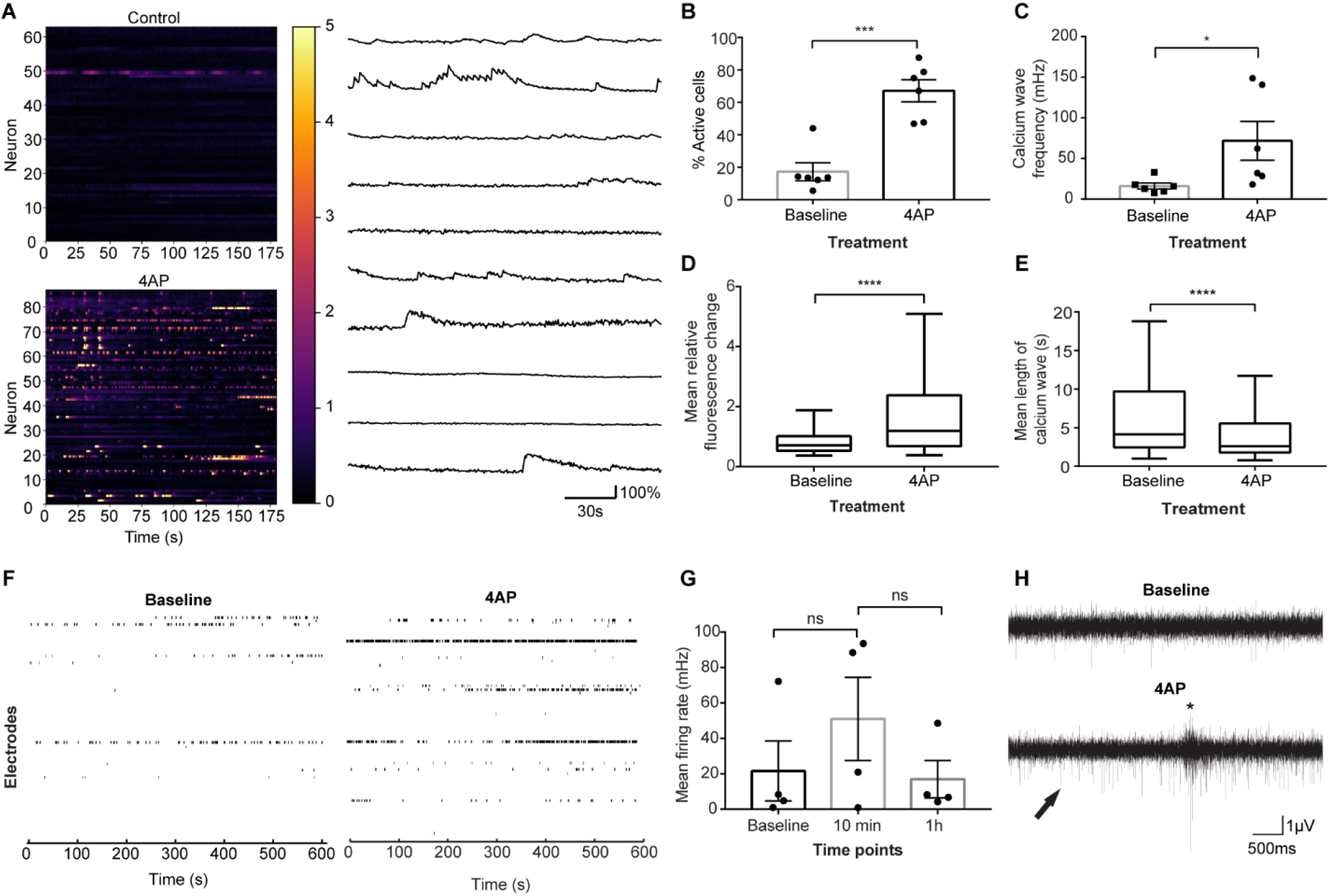
Effect of 4AP on neuronal activity in 90-day-old cerebral spheroids. (A) *Left:* Heatmap of calcium-associated fluorescence over time in a spheroid before exposure to 4AP (upper) and after exposure to 4AP (lower). Each line represents a different cell and the color bar represents the fold-change in calcium-associated fluorescence over each cell’s baseline. *Right:* Individual traces of calcium-associated fluorescence over time of representative cells (B) Proportion of active cells before and after treatment with 4AP. Student’s *t*-test, *p* = 0.0003. Bars represent mean +/− SEM. Each dot represents an individual spheroid. *** *p* < 0.001. (C) Calcium wave frequency in active cells exposed or not to 4AP. Wilcoxon Rank Sum test, *p* = 0.0152. Bars represent mean +/− SEM. Each dot represents an individual spheroid. * *p* < 0.05. (D) Mean calcium transient amplitude. The mean calcium transient amplitude was calculated for each cell from 6 individual spheroids for each condition (n = 282 cells in 4AP group and 95 cells in Baseline group. Wilcoxon Rank Sum test, *p* < 0.0001. The line represents the median, sides of the box represent the upper and lower quartile and whiskers represent maximum and minimum values. **** *p* < 0.0001. (E) Mean calcium transient duration. The mean calcium transient duration was calculated for each cell from 6 individual spheroids for each condition (n = 282 cells in 4AP group and 95 cells in Baseline group. Wilcoxon Rank Sum test, *p* < 0.0001. The line represents the median, sides of the box represent the upper and lower quartile and whiskers represent maximum and minimum values. **** *p* < 0.0001. (F) Representative raster plot of neuronal spikes before (*left*) and after (*right*) exposure to 100μM 4AP. Each line represents an individual electrode. (G) Mean firing rate upon exposure to 4AP. ANOVA repeated measures with Holm Sidak’s correction for multiple comparisons, F_(1,439, 4,316)_ = 0.6389 *p* = 0.5226. Each dot represents an individual spheroid. Bars represent mean +/− SEM. ns: not significant. (H) Example traces of local field potentials recorded before (upper) and after (lower) 10-minute exposure to 4AP showing an increase in spikes (arrow) as well as the presence of bursts (*). For graphs, each dot represents an individual spheroid.

**Figure 4:**
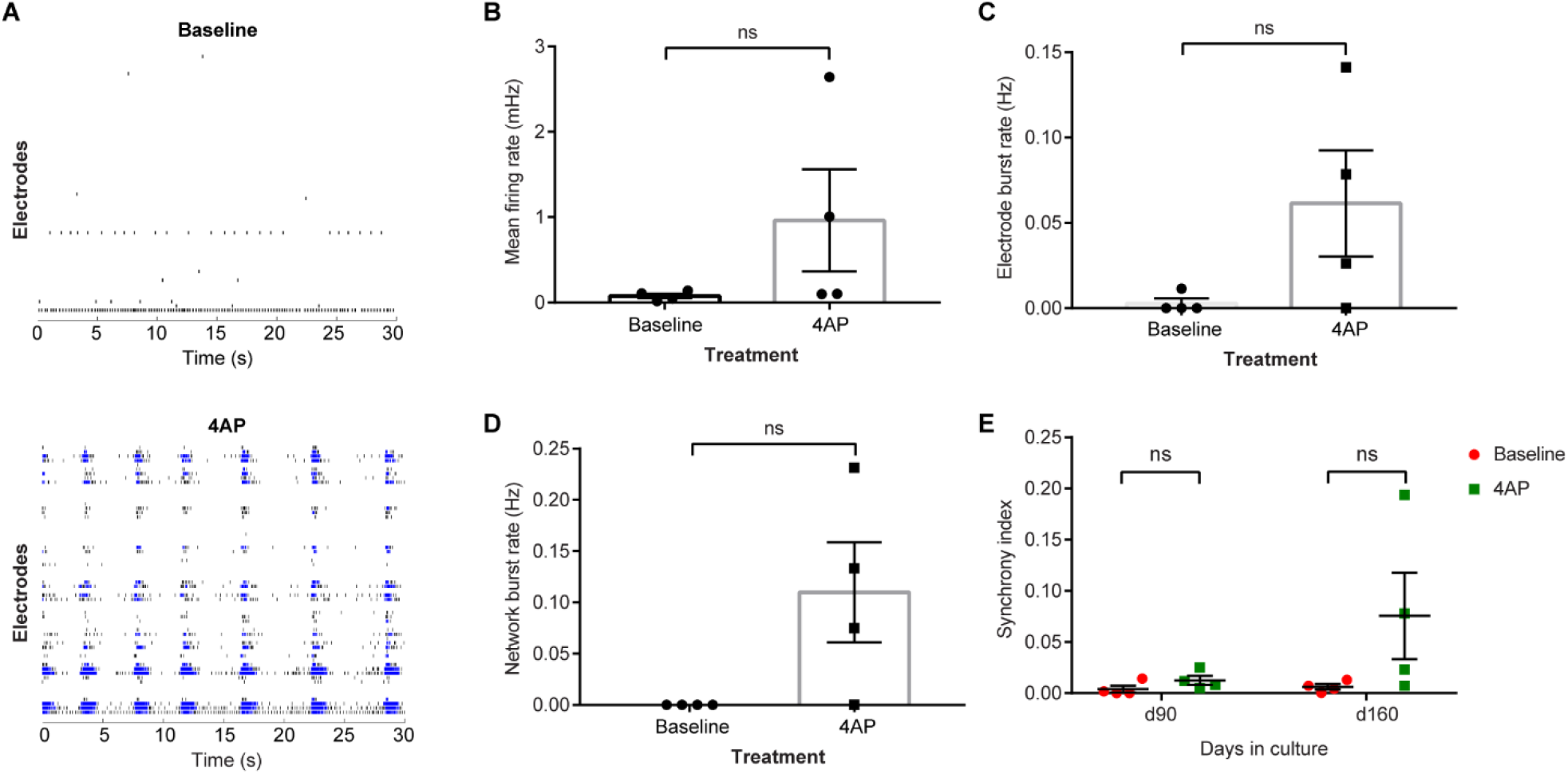
Effect of 4AP on neuronal activity in 160-day-old cerebral spheroids. (A) Representative raster plots of a 160-day-old cerebral spheroids before (*upper*) and after exposure (*lower*) to 4AP. Exposure to 4AP triggered network bursts characterized by synchronous bursting of several electrodes. Black marks indicate individual spikes while blue marks indicate electrode bursts. (B) Mean firing rate in spheroids before and after exposure to 4AP. Paired Student’s *t*-test, *p* = 0.2327. Bars represent mean +/− SEM. Each dot represents an individual spheroid. ns: not significant. (C) Electrode burst frequency in spheroids before and after exposure to 4AP. Paired Student’s *t*-test, *p* = 0.1750. Bars represent mean +/− SEM. Each dot represents an individual spheroid. ns: not significant. (D) Network burst frequency in spheroids before and after exposure to 4AP. Paired Student’s *t*-test, *p* = 0.1097. Bars represent mean +/− SEM. Each dot represents an individual spheroid. ns: not significant. (E) Synchrony index before and after exposure to 4AP at day 90 and day 160. Paired Student’s *t*-test, *p _day 90_* = 0.1762 and *p _day 160_* = 0.1528. Bars represent mean +/− SEM. Each dot represents an individual spheroid. ns: not significant.

These findings reveal that synchronized network bursts could be triggered in cerebral spheroids with 4AP after 160 days of culture. This suggests that while 4AP increases neuronal activity in 90-day-old spheroids, these spheroids require a longer time in culture to generate synchronized activity.

### Activity-induced neurogenesis in cortical spheroids

Activity-induced neurogenesis has been documented both *in vitro* and *in vivo*. We next examined whether neurogenesis was similarly modulated in spheroids by increasing neuronal activity with 4AP. We evaluated neurogenesis in 90-day-old spheroids as a significant number of SOX2+ neural stem cells are still present and organized in neural tube-like structures at that stage compared to older spheroids. First, we analyzed the effect of 4AP on proliferation and noted an approximately 2-fold decrease in the number of proliferative Ki67-positive cells after 24 h of 4AP exposure (2.2963% vs 4.367%; Figure 5A). We next identified a small but not significant increase in apoptotic cleaved-caspase3-positive cells in 4AP treated organoids (0.1875% vs 0.3019%; Figure 5B).

**Figure 5:**
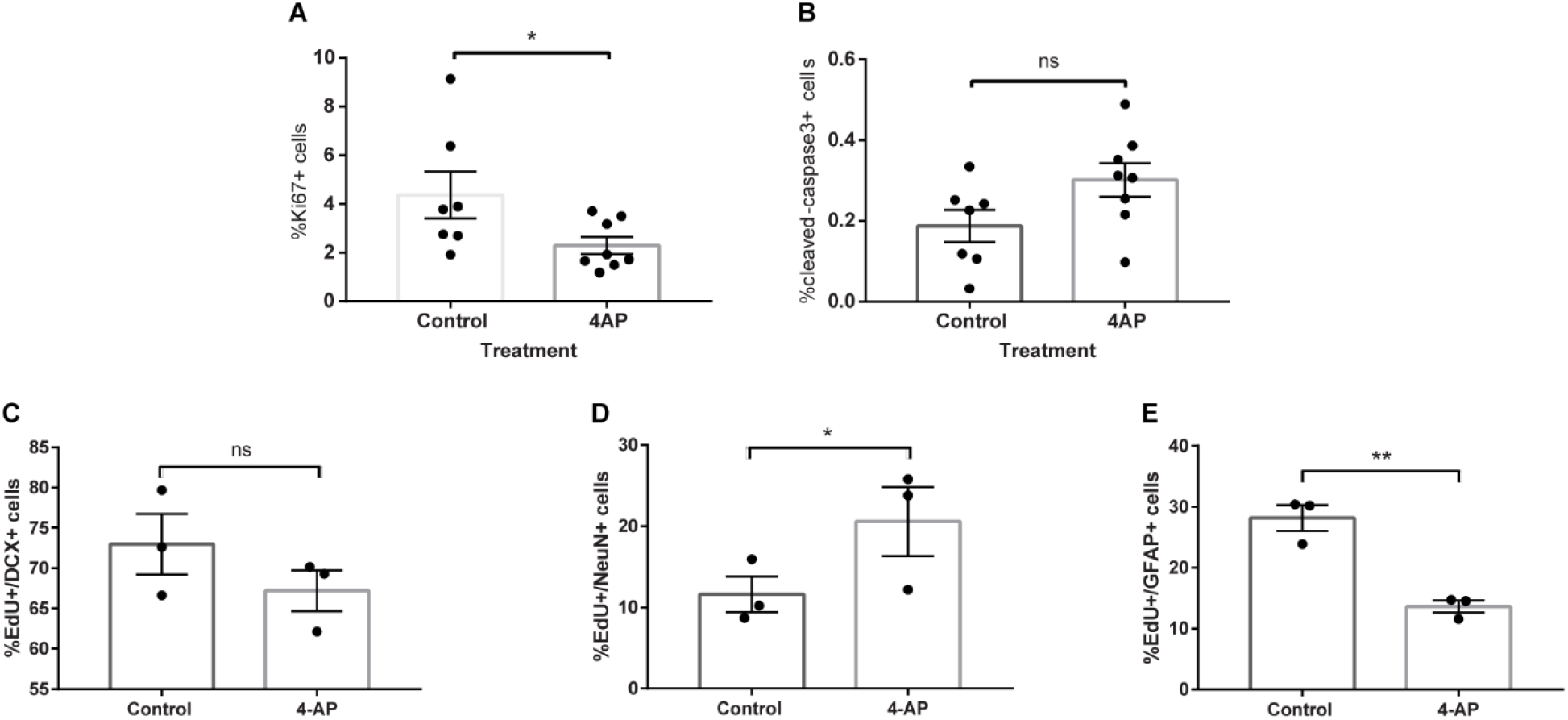
Activity-induced apoptosis, proliferation, neurogenesis and gliogenesis in cerebral spheroids. (A) Proportion of Ki67-positive proliferative cells 24 h after treatment with 4AP or vehicle (water). Student’s *t* test with Holm Sidak’s correction for multiple comparisons, *p* = 0.0331. Bars represent mean +/− SEM. Each dot represents an individual spheroid. \**p* < 0.05. (B) Proportion of cleaved-caspase 3-positive apoptotic cells 24h after treatment with 4AP or vehicle (water). Student’s *t* test with Holm Sidak’s correction for multiple comparisons, *p* = 0.0995. Bars represent mean +/− SEM. Each dot represents an individual spheroid. ns: not significant. (C) Proportion of EdU+/DCX+ cells in EdU+ cells. Student’s *t* test with Holm Sidak’s correction for multiple comparisons, *p* = 0.0880. Bars represent mean +/− SEM. Each dot represents an individual spheroid. ns: not significant. (D) Proportion of EdU+/NeuN+ cells in EdU+ cells. Student’s *t* test with Holm Sidak’s correction for multiple comparisons, *p* = 0.0331. Bars represent mean +/− SEM. Each dot represents an individual spheroid. \**p* < 0.05. (E) Proportion of EdU+/GFAP+ cells in EdU+ cells. Student’s *t* test with Holm Sidak’s correction for multiple comparisons, *p* = 0.0073. Bars represent mean +/− SEM. Each dot represents an individual spheroid. \*\**p* < 0.01.

We then investigated whether 4AP affected the differentiation of NSCs by tagging proliferating cells with EdU just prior to stimulation with 4AP and assessing their differentiation 2 weeks later using antibodies against immature neurons (DCX), mature neurons (NeuN) or glial cells (GFAP). A higher proportion EdU-positive cells in 4AP-treated spheroids expressed the mature neuron marker NeuN compared to control spheroids (20.6% vs 11.6% respectively). No statistically significant difference was found in the proportion of EdU+/DCX+ cells between 4AP-treated and control spheroids (47.4% vs 58.3% respectively). In contrast, a higher proportion of EdU-positive cells expressed the glial cell marker GFAP in control spheroids compared to 4AP-treated spheroids (28.2% vs 13.6% respectively; Figure 5C-E). Overall, this suggests that proliferating cells at the time of 4AP exposure are more likely to differentiate into neurons and less into glial cells under these conditions.

### Activity-dependent gene expression in spheroids

To investigate whether 4AP triggered activity-dependent gene expression in neurons within spheroids, we evaluated the proportion of postmitotic NeuN+ neurons that expressed the immediate early gene cFos in organoids exposed to 4AP compared to controls. These NeuN+/cFos+ neurons represent putative recently stimulated neurons. As expected, the proportion of NeuN+/cFos+ double positive neurons was higher in spheroids exposed to 4AP than control spheroids (51.1% vs 15.5% respectively; Figure 6A). This result demonstrates that 4AP increased neuronal activity and induced a higher expression of the IEG cFOS in post-mitotic neurons.

**Figure 6:**
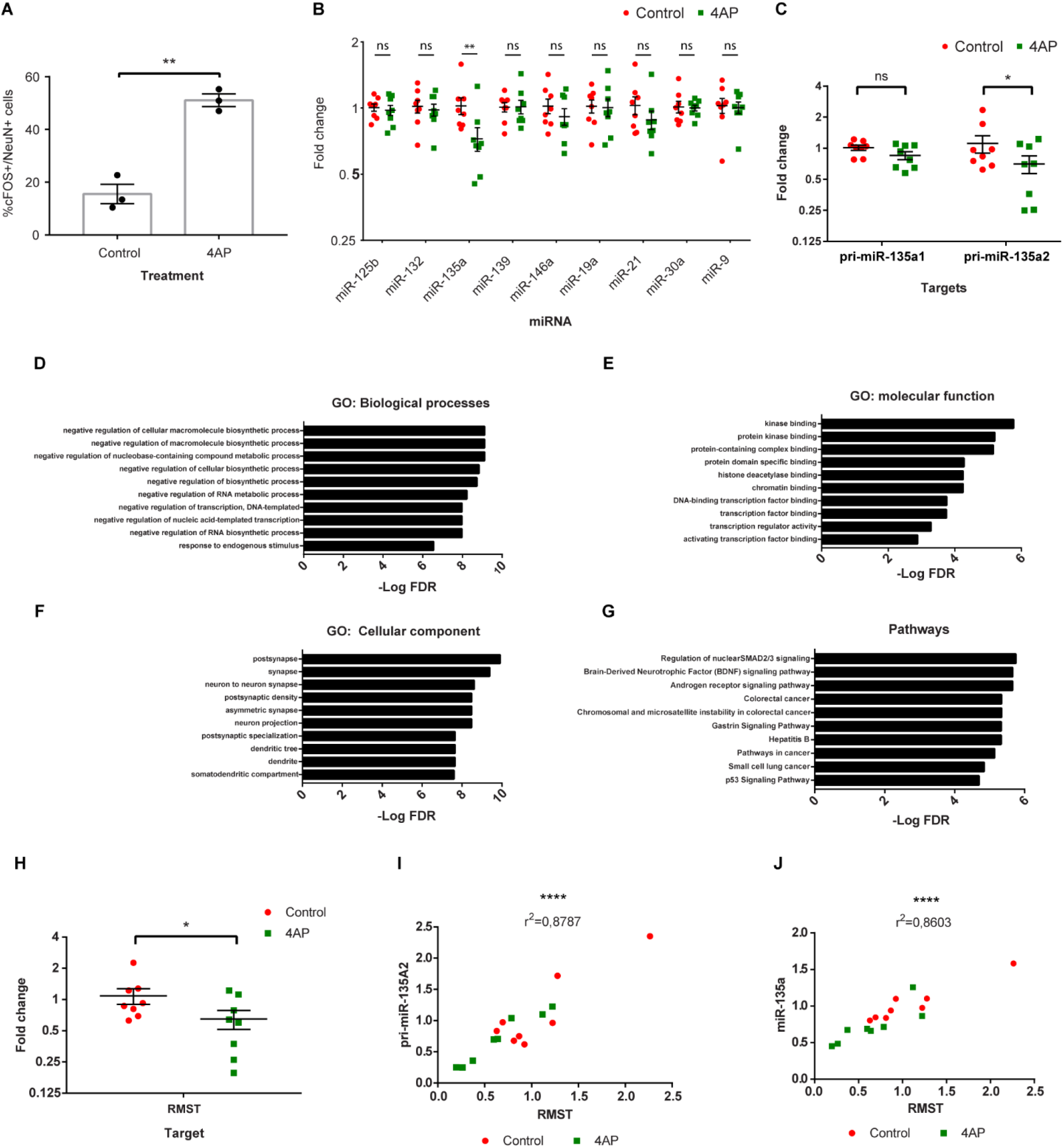
Activity-induced expression of cFOS, epilepsy-associated miRNAs and RMST in cerebral spheroids. (A) Proportion of cFOS+/NeuN+ in NeuN+ neurons in organoids exposed or not to 4AP. Student’s *t*-test, *p* = 0.0034. Bars represent mean +/− SEM. Each dot represents an individual spheroid. \*\**p* < 0.01. (B) Expression of epilepsy-associated miRNAs in spheroids. ANOVA followed by multiple Student’s *t-* test with Holm Sidak’s test for multiple comparisons. F _(1, 126)_ = 5.63, *p* = 0.0192, *p_miR-135a_* = 0.0053. Bars represent mean +/− SEM. Each dot represents an individual spheroid. N=8 spheroids in each group. ns: not significant, \*\**p* < 0.01. (C) Expression of pri-mir-135a1 and pri-mir-135a2 in cerebral spheroids exposed to 4AP or not. Expression was measured with RT-qPCR. ANOVA followed by multiple Student’s *t-*test with Holm Sidak’s test for multiple comparisons. F _(1, 28)_ = 5.401, *p* = 0.0276, *p _pri-miR-135a2_* = 0.0455. Bars represent mean +/− SEM. Each dot represents an individual spheroid. ns: not significant, \**p* < 0.05. (D), (E), (F), (G) Enrichment analysis for miR-135a targets. Top 10 terms for GO biological processes (D), GO molecular function (E), GO cellular components (F) and KEGG pathway (G). FDR = False Discovery Rate. (H) Expression of RMST in cerebral spheroids exposed to 4AP or not. Expression measured with RT-qPCR. Student’s *t*-test, *p* = 0.0453. Bars represent mean +/− SEM. Each dot represents an individual spheroid. \**p* < 0.05. (I) and (J) Correlation between RMST expression and pri-mir-135A2 and miR-135a respectively. Pearson correlation coefficient. Each dot represents an individual spheroid. \*\*\*\**p* < 0.0001.

MicroRNAs are involved in the regulation of gene expression and implicated in neuronal differentiation of NSCs (Olde Loohuis et al., 2012; Shen and Temple, 2009). Moreover, the expression of specific miRNAs is induced by neuronal activity (Eacker et al., 2011). We therefore hypothesized that, if certain miRNAs were implicated in activity-induced neurogenesis, we would observe differential expression of these miRNAs in spheroids treated with 4AP. We focused on 9 miRNAs previously demonstrated to participate in neurogenesis and consistently differentially expressed in rodent models of epilepsy as well as human epileptic tissue (Korotkov et al., 2017). Of the 9 miRNAs selected, only miR-135a was differentially expressed between treated and control spheroids; its expression was significantly decreased in spheroids exposed to 4AP (Fold change: 0.72; Figure 6B). To identify the potential functional significance of miR-135a differential expression after 4AP treatment, we investigated the 512 mRNA targets of miR-135a reported by one or more databases of validated miRNA-mRNA interactions. These mRNA targets were subjected to GO and KEGG enrichment analysis to identify potential functional pathways. GO enrichment analysis for biological processes revealed a significant enrichment for terms related to negative regulation of biosynthetic processes and transcription. GO: molecular function terms were enriched for kinase binding as well as transcription factors and chromatin binding factors. GO: cellular components were enriched for neuronal components such as the synapse and dendrites. Finally, KEGG pathway terms were enriched for SMAD2/3 and BDNF signaling suggesting a role of miR-135a targets in growth and differentiation (Figure 6D-G). This supports roles for miR-135a in activity-induced neurogenesis at least in part by targeting transcription factors that, once activated, drive NSCs toward the neuronal lineage.

The decrease in miR-135a expression after exposure to 4AP could be either due to decreased transcription or increased turnover. To investigate this further, we measured the expression of pri-miR-135a by RT-qPCR. MiR135a is encoded by two genes: *MIR135A1* on human chromosome 3 and *MIR135A2* on human chromosome 12. We designed primers specific for each stem-loop transcript (pri-miR-135a1, pri-miR-135a2) and measured the expression of each transcript using RT-qPCR. Interestingly, only pri-miR-135a2 showed a decrease in expression in the spheroids treated with 4AP (Figure 6C). This suggests that the observed decrease in mature miR135a is due to at least in part to a decrease in transcription of the *MIR135A2* gene or increased turnover of pri-miR-135a2 prior to processing into its mature form.

The *MIR135A2* gene is located in the last intron of *RMST*, a gene coding for the long non-coding RNA RMST (Anderegg and Awatramani, 2015). The transcription of intragenic miRNAs can be either under the control of promoters of the host gene in which they are located, or through nearby independent promoters. To begin to elucidate whether *MIR135A2* transcription is under the control of the *RMST* promoter in our spheroid model, we measured the expression of the RMST transcript in spheroids treated with 4AP and in control spheroids and observed a decrease in RMST expression in 4AP-treated spheroids (0.6-fold change; Figure 6H). We then plotted the level of expression of pri-mir-135a2 and mir-135a as a function of the level of expression of RMST and observed a strong correlation between them (r^2^ = 0.8787 and r^2^ = 0.8603 respectively; Figure 6I-J). Altogether, this suggests that the decrease in miR-135a expression is due at least partly to a decrease in *RMST* transcription, although increased turn-over of the RMST primary transcript may also contribute to the decrease in pri-mir135a2 levels.

## DISCUSSION

Here we present some of the first evidence to support activity-induced neurogenesis and concomitant changes in miRNA expression in cerebral spheroids. 4AP increased neuronal activity in 90-day-old spheroids although longer time in culture was required for the appearance of synchronized activity This is in line with a previous study that has reported little synchronization in 2-4 months old organoids but a gradual increase in neuronal spontaneous depolarization events until 8 months of culture (Trujillo et al., 2019). The increase in newly formed neurons (EdU+/NeuN+) and decrease in newly formed glial cells (EdU+/GFAP+) in spheroids treated with 4AP constitutes striking evidence for activity-induced neurogenesis in spheroids. Of note, similar evidence highlighting the effect of neuronal activity on neuronal and glial differentiation has been described both *in vitro* and *in vivo* (Deisseroth et al., 2004; Ma et al., 2009; Song et al., 2016; Tozuka et al., 2005). However, we did not determine whether the increase in neuronal differentiation observed in spheroids is mediated directly through the increase in neuronal activity by 4AP or through an independent action of 4AP on NSCs. 4AP blocks potassium channels leading to an increase in neurotransmitter release in pre-synaptic neurons and inhibits repolarization in post-synaptic neurons, but other effects notably on NSCs have not been studied (Avoli and Jefferys, 2016).

The mechanisms underlying activity-induced neurogenesis remain unknown. Our data hints at the potential implication of miR-135a. This miRNA was shown to be important in exercise-induced neurogenesis in the mouse hippocampus where its downregulation was necessary for the formation of new neurons (Pons-Espinal et al., 2019). We found that the expression of miR-135a and its precursor pri-mir-135a2 is correlated with the expression of RMST. As the *MIR135A2* gene is located in the last intron of *RMST*, this would suggest a common promoter. RMST was shown to interact with SOX2 and its binding to the promoters of neurogenic genes, inducing neuronal differentiation of embryonic cells (Ng et al., 2013). However, in this study, suppressing *RMST* expression did not decrease the expression of miR-135a2. It is possible that the RMST transcript is quickly spliced and pri-miR-135a2 rapidly generated after transcription. Therefore, the RMST mRNA is more abundant after splicing and preferentially targeted by siRNA. If this is the case, siRNA-mediated knock down of RMST would not be expected to significantly affect miR-135a levels. *RMST* expression is repressed by REST which has been shown to inhibit the expression of several neuronal genes and prevent neuronal differentiation (Conaco et al., 2006; Ng et al., 2013). It is unknown whether the decrease in *RMST* expression in our study is due to increased REST binding or not. Moreover, *RMST* transcription is at least partly under the control of the transcription factor Lmx1b and activated during Wnt signalling, which promotes neurogenesis. It has been shown that miR-135a2 in turn targets the transcripts of several Wnt pathway effectors, thereby acting as a negative feedback regulator to constrain neurogenesis (Anderegg and Awatramani, 2015). Overall, a decrease in miR-135a expression may represent a permissive event for activity-induced neurogenesis and several other yet to be identified factors are likely to play additional roles.

Another question that remains unanswered is whether the decrease in miR-135a expression occurs specifically in NSCs or in other cell types, which then indirectly influence neuronal differentiation of NSCs. It has been shown that astrocytes can express miR-135a which participates in neuronal survival (Chu et al., 2016). Evaluating the expression of miR-135a in individual cell types using fluorescent *in situ* hybridization or cell-sorting prior to RT-qPCR would be necessary to specifically address that question.

In conclusion, we have examined effects of 4AP on cerebral spheroids. 4AP enhanced neuronal differentiation and decreased glial differentiation of neural progenitors in this model. This is the first evidence of activity-induced neurogenesis in human cerebral spheroids. We identified miR-135a as one potential mediator of activity-induced neurogenesis, although the mechanisms underlying its function remain undetermined.

## EXPERIMENTAL PROCEDURES

### Cerebral spheroid generation

Human iPSCs were generated from neurologically normal individuals as described previously (Zimmerman et al., 2020). Briefly, iPSCs were generated from dermal skin punch (3mm^3^)-derived fibroblasts obtained with informed written consent from healthy adult control subjects at the Massachusetts General Hospital (MGH), Department of Psychiatry, and reprogrammed as iPSCs using a nonintegrative, mRNA-based technology (Cellular Reprogramming, Inc. (www.cellular-reprogramming.com)). The study protocol was approved by the Institutional Review Board of the Massachusetts General Hospital in accordance with U.S. Common Rule ethical guidelines. Human iPSCs) were cultured in Essential 8 medium (ThermoFisher, Waltham, MA USA) or Nutristem hPSC XF medium (Biological Industries, Kibbutz Beit-Haemek, Israel) plated on Matrigel (Corning, Glendale, AZ USA) or Laminin-521 (Stemcell technologies, Vancouver, BC Canada) depending on the cell line. Cortical spheroids were generated from iPSCs following a previously described protocol with slight modifications. iPSCs were dissociated into single cells using Accutase (Gibco ThermoFisher). Cells were resuspended in Essential 8 medium supplemented with Y26732 ROCK inhibitor (Selleckchem, Houston, TX USA) at 10μM and 10,000 cells were added to each well of an ultra-low attachment, U-bottom 96-well plate (ThermoFisher) to reaggregate overnight. The next day (noted as D0), medium was changed for E6 (Stemcell technologies) supplemented with 10μM SMAD inhibitor SB431542 (Tocris, Bristol UK) and 2.5 μM SMAD inhibitor Dorsomorphin (Sigma-Aldrich, St. Louis, MO USA). This medium was replaced every day for 5 days. On day 6, spheroids were transferred to an ultra-low attachment 6-well plate placed on an orbital shaker at 65 rpm and medium was changed to Neurobasal medium (Gibco ThermoFisher) supplemented with 2% B27 supplements (Gibco ThermoFisher), 1% Penicillin/Streptomycin 10000U/L (Gibco ThermoFisher) and 1% GlutaMax (Gibco ThermoFisher). Basic Fibroblast Growth Factor (bFGF; 20 ng/ml, Stemcell technologies) and Epidermal Growth Factor (EGF; 20ng/ml, Stemcell technologies) were added from day 6 to day 25. Medium was changed every day until day 15 and then every other day until day 25. On day 25, bFGF and EGF were replaced with Brain-derived Neurotrophic Factor (BDNF; 20ng/ml, Stemcell technologies) and Neurotrophin-3 (NT-3; 20 ng/ml, Stemcell technologies) and the culture medium was changed every 3-4 days. On day 43, BDNF and NT-3 supplementation was stopped, and medium was changed every 3-4 days until spheroids were collected (see supplemental figure 1). Use of the patient-derived iPSC lines and generation of cerebral organoids was approved by the University of Guelph Research Ethics Board (REB 17– 11–012). Three different iPSC lines were used from 3 different individuals.

### Immunofluorescence

At selected time points, spheroids were collected, fixed in 4% paraformaldehyde (PFA), dehydrated in a 30% sucrose solution and 20 μM cryosections were acquired with a cryostat. Sections were permeabilized and blocked with 1% Bovine Serum Albumin (BSA, Sigma-Aldrich) and 0.3% Triton-X (FisherScientific) in PBS for 1 hour at room temperature before incubation with primary antibodies overnight followed by incubation with Alexa Fluor 488 or 594-conjugated secondary antibodies (Supplementary Table 1) for 2 h at room temperature. Cells were then counterstained with 4′,6-diamidino-2-phenylindole (DAPI). For Ki67 immunohistochemistry, a heat-mediated antigen retrieval step was performed by incubating the slide in a sodium citrate buffer (10 mM sodium citrate, 0.05% Tween-20, pH = 6) for 20 minutes in a water bath at 95°C prior to primary antibody incubation. Z-stacks were acquired on a FV1000 Olympus confocal microscope running Olympus Fluoview software version 4.3 (Olympus, Tokyo, Japan). All stacks were acquired with the same settings between different biological replicates. Z-stacks and multi channel images were reconstructed using ImageJ (version 1.52e, Rasband, W.S., ImageJ, U. S. National Institutes of Health, Bethesda, Maryland, USA, https://imagej.nih.gov/ij/, 1997-2018). Cells expressing each antigen were manually counted using ImageJ and averaged over 3 different regions of each organoid. The total number of cells was estimated by counting the numbers of DAPI stained nuclei using the Analyze Particle ImageJ plugin. Negative controls were performed by omitting the primary antibodies to control for non-specific binding of the secondary antibodies.

### EdU labelling

The EdU staining proliferation kit was used according to manufacturer’s recommendations (Abcam). Briefly, spheroids were incubated with 20 μM EdU for 2 hours before being exposed to 4AP or vehicle (distilled water). Two weeks later, spheroids were fixed, dehydrated and cryosectioned. Sections were then permeabilized and EdU was tagged using the fluorescent azide i647 before being incubated with primary and secondary antibodies as described above. Images were acquired and processed as described above.

### Electrophysiology

For whole-cell recording, spheroids were collected at selected time points and embedded in 3% Agarose. Embedded organoids were then cut in 400 μm thick slices with a Leica VT 1200 vibrating microtome (Leica Biosystems, Concord, Ontario, Canada) and slices were transferred into artificial cerebrospinal fluid (aCSF:128 mM NaCl, 10 mM D-glucose, 26 mM NaHCO_3_, 2 mM CaCl_2_, 2 mM MgSO_4_, 3 mM KCl, 1.25 mM NaH_2_PO_4_, pH = 7.4) bubbled with 95% O_2_ and 5% CO_2_ at 30°C for 1 hour to recover. Slices were then transferred to a recording chamber (Warner Instruments, Hamden, CT USA) mounted on the stage of an Axioskop FS2 Microscope (Carl Zeiss Canada, Toronto, Ontario, Canada). Slices were superfused continuously with aCSF bubbled with 95% O_2_ and 5% CO_2_ at room temperature. Spheroids smaller than 2 mm were not sliced and recorded as intact spheroids. Whole-cell recordings of putative neurons were performed with borosilicate glass pipette electrodes filled with an internal solution containing 120 mM K-gluconate, 5 mM KCl, 2 mM MgCl_2_, 4 mM K2-ATP, 400 μM Na_2_-GTP, 10 mM Na_2_-phosphocreatine and 10 mM HEPES buffer (adjusted to pH 7.3 with KOH). Pipette resistance was 4-6 MΩ. Whole-cell recordings were acquired at 20 kHz and lowpass filtered at 2 kHz using a Multiclamp 700B amplifier and Digidata 1440A digitizer (Molecular Devices, San Jose, CA, USA). Neuronal intrinsic excitability (input/output curve, rheobase) was assessed in current clamp mode by injecting positive current steps for 500 ms each. The frequency of spontaneous action potentials was measured in current clamp mode at resting membrane potential. The frequency and amplitude of spontaneous excitatory post-synaptic currents (sEPSCs) were recorded in voltage clamp mode with cells held at −75 mV.

For multielectrode array (MEA) recordings, a single whole spheroid was transferred just prior to recording into a well of a 6-well MEA with 64 low-impedance platinum microelectrodes (Axion Biosystems, Atlanta, GA USA) previously coated with 100 μg/ml poly-d-lysine (Sigma-Aldrich) and 10 μg/ml Laminin (Sigma-Aldrich). Before recording, culture medium was aspirated so that a minimal amount of medium (approximately 200 μL) covered the spheroid to minimize detachment and movement of spheroids. Recordings were performed on a Maestro Edge 384 channel system (Axion Biosystems) and AxIS software Spontaneous Neural Configuration (Axion Biosystems). Recordings were performed in culture medium at 37°C with 5% CO2 for 10 minutes. Baseline recordings were obtained after which spheroids were incubated with 4AP at 100 μM (Sigma-Aldrich). This dose of 4AP was chosen based on previous publications (Gonzalez-Sulser et al., 2011; Heuzeroth et al., 2019; Hsiao et al., 2015). Analysis of the MEA recordings was performed with the Axion Biosystems Neural Metrics Tool using MATLAB scripts (version R2021a). Spikes were detected using an adaptative threshold of 5.5 times the standard deviation of the estimated noise for each electrode. Electrode bursts were defined as a minimum of 5 spikes with a maximum inter-spike interval of 100 ms. A network burst was defined as a minimum of 10 spikes under a maximum inter-spike interval of 100 ms with a minimum of 20% active electrodes (Trujillo et al., 2019).

### Calcium imaging

Whole organoids were incubated for 30 minutes in culture medium with 2 μM of Fluo4-AM. Organoids were then washed and imaged on a FV1000 Olympus confocal microscope running Olympus Fluoview software version 4.3 (Olympus, Tokyo, Japan) using a 488nm excitation laser. Images were acquired every 0.5 s for 3 minutes. After acquiring baseline recordings, organoids were stimulated with 100 μM of 4AP. The fluorescence of individual neurons was followed over time by manually drawing a region-of-interest (ROI) around each neuron and measuring the mean ROI fluorescence for each frame using ImageJ. Background fluorescence was subtracted from each measure and the relative change in fluorescence of each neuron was expressed as (F-F0)/F0 where F represents the mean fluorescence of a neuron at a particular time and F0 represents the minimum fluorescence of that neuron. Calcium peaks were identified using custom Python scripts (version 3.8, Python Software Foundation, http://www.python.org) to identify local maxima. Only peaks with a prominence of 0.3 (representing an increase of 30% in fluorescence compared to the nearest local minima) were kept, discarding peaks due to noise only. Traces were visually inspected to confirm automatic peak detection. The calcium peak frequency, number of calcium peaks per cell and the number of cells showing at least one peak (active cells) were counted. The mean amplitude and duration of calcium transients for each active cell was also recorded.

### RT-qPCR

Total RNA of individual spheroids was isolated with the miRNeasy micro Kit (Qiagen, Hilden, Germany) according to manufacturer’s instructions. An on-column DNase digestion was performed with the RNase-free DNase set (Qiagen). RNA was quantified with the Nanodrop 2000c (ThermoFisher). For both mRNA and miRNA, 500 μg of total RNA was reverse transcribed. Messenger RNAs were reverse transcribed with qScript complementary DNA (cDNA) SuperMix (Quantabio, Berverly, MA USA). MicroRNAs were first polyadenylated and then reverse transcribed using the qScript miRNA cDNA Synthesis kit (Quantabio). Real-Time quantitative Polymerase Chain Reaction (RT-qPCR) was performed using a CFX96 Touch Real-Time PCR Detection System (Bio-Rad Laboratories) using 3 ng of cDNA per reaction and Sensifast SYBR No-ROX mix (Bioline Corporation, Alvinston, ON, Canada). Complementary DNA of mRNA was amplified using a pair of specific forward and reverse primers (Supplementary Table 2). MicroRNAs were amplified using a specific forward primer and PerfeCTa Universal PCR primer (Quantabio). Standard curves were performed for each primer pair to calculate primer efficiency. Primer specificity was assessed by performing a melting curve, examining the melting curve for multiple peaks, and separating the PCR products on a 1% Agarose gel and assessing the products for multiple bands. The stability of reference gene expression was evaluated using the GeNorm method (Vandesompele et al., 2002). GAPDH and HPRT were utilized as references for mRNA targets while miR-17 and miR-181c were utilized for miRNA normalization.

### MicroRNA target functional annotation enrichment analysis

Validated mRNA targets of differentially expressed miRNAs were retrieved from the databases miRecords, miRTarbase v8 and Tarbase v8 (Huang et al., 2019; Sethupathy, 2005; Xiao et al., 2009). Enrichment for specific biological processes, molecular function and cellular compartment GO terms, as well as KEGG pathways, were analyzed with the ToppFunn module of the ToppGene suite using a probability density function with Benjamini-Hochberg False Discovery Rate (FDR) correction (Chen et al., 2009).

### Statistics

Graphing and statistical analyses were performed with GraphPad Prism 7 (GraphPad Software, La Jolla, CA USA). Normal distribution of data was verified with a Shapiro Wilk Test. If the data was normally distributed, differences between means were analyzed by Student’s two-tailed *t*-test or One-way ANOVA followed by multiple Student’s *t-test* with Holm Sidak’s correction for multiple comparisons when there were more than 2 groups. If the data was not normally distributed, a Wilcoxon Rank Sum test was performed. For proportions, a logit-transformation was performed and Student’s *t*-tests were used on the logit transformed-data. Gene expression data from RT-qPCR was calculated using the ΔΔCT method using qbase+ software, version 3.2 (Biogazelle, Zwijnaarde, Belgium - www.qbaseplus.com) and log_2_ transformed to assume a normal distribution (Pfaffl, 2001). Correlations between gene expression were performed using a Pearson correlation. The number and nature of replicates is indicated in each figure. A *p*-value < 0.05 was considered statistically significant. Data shown represent the mean +/− standard error of the mean (SEM). No prior sample size calculation was performed.

### Code availability

Code and data are available upon request.

## ACKNOWLEDGMENTS

This work was supported by grants from: The OVC Pet Trust Fund and NSERC (to J LaMarre).

## AUTHOR CONTRIBUTIONS

T.P. and J LaMarre designed the project. T.P., E. H. and N. W. performed all experiments. T.P. and J. LaMarre analyzed the data. F. J., C. B., S. D. S., R. H. P., M. P., L. G., J. Lalonde, and J. LaMarre provided resources, conceptualization, and intellectual content. T. P. and J. LaMarre wrote the paper with feedback from all authors.

---

**Supplementary Figure 1:**
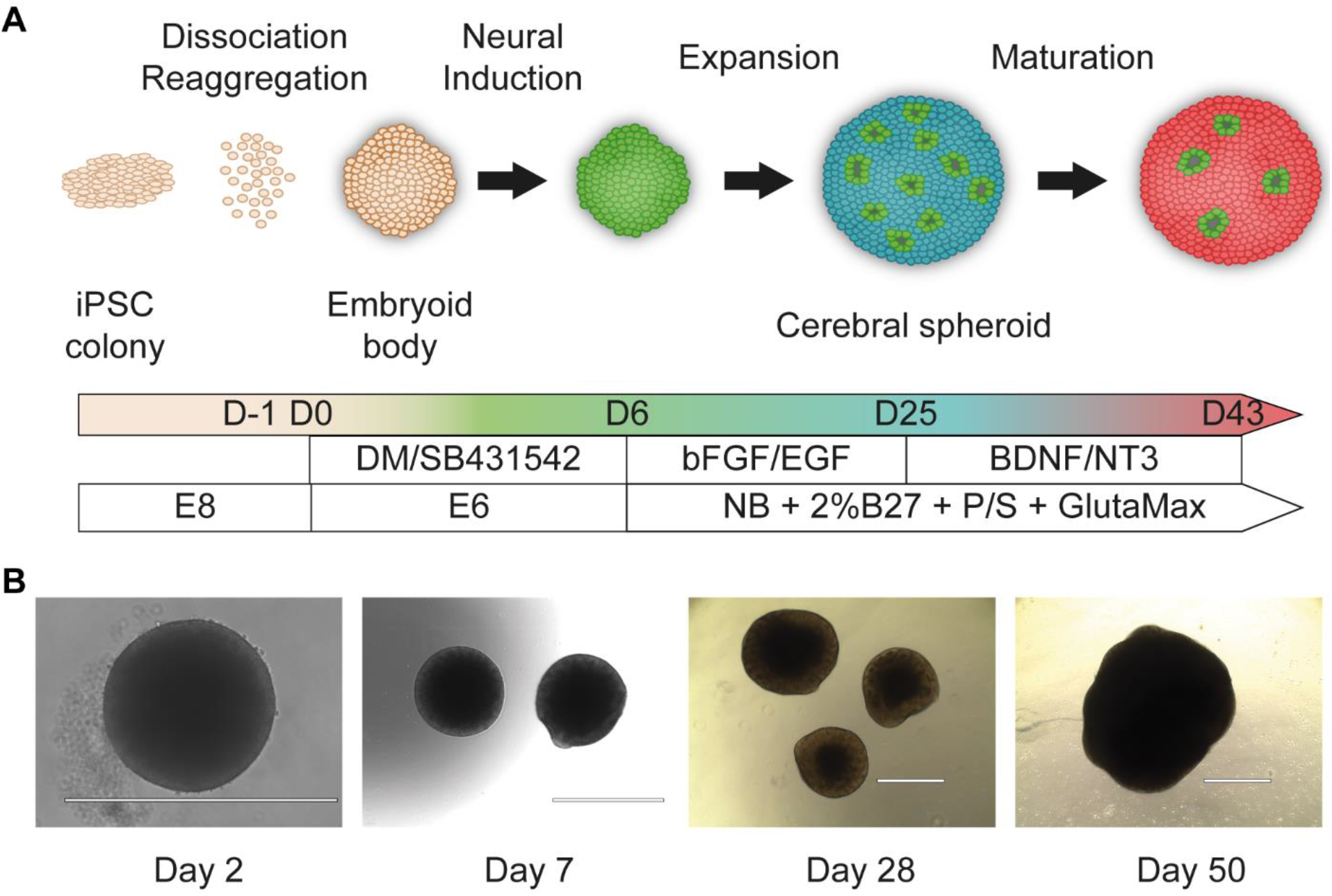
Overview of cerebral spheroid development. (A) Schematic diagram of cerebral spheroid protocol. Adapted from Yoon et al., 2019. (B) Representative bright field images over time in culture. Bar = 1 mm BDNF: brain-derived neurotrophic factor, bFGF: basic fibroblast growth factor, DM: dorsomorphin, EGF: epidermal growth factor, iPSC: induced pluripotent stem cells, NB: Neurobasal medium, NT3: neurotrophin 3, P/S: Penicillin/Streptomycin

