## Supplementary information for "Human Cerebral Spheroids Undergo Activity Dependent Changes In Cellular Composition And Microrna Expression"

### Supplemental information

***Supplementary Figure 1:*** **Overview of cerebral spheroid development**

(A) Schematic diagram of cerebral spheroid protocol. Adapted from Yoon et al., 2019.

(B) Representative bright field images over time in culture. Bar = 1 mm

BDNF: brain-derived neurotrophic factor, bFGF: basic fibroblast growth factor, DM: dorsomorphin, EGF: epidermal growth factor, iPSC: induced pluripotent stem cells, NB: Neurobasal medium, NT3: neurotrophin 3, P/S: Penicillin/Streptomycin

**<u>Supplementary Table 1: Primary and secondary antibodies used</u>**

| **Antigen** | **Host species** | **Working dilution** | **Manufacturer (catalog number)** |
|---|---|---|---|
| Sox2 | Rabbit | 1:500 | Abcam (ab97959 |
| Nestin | Rabbit | 1:500 | Abcam (ab92391) |
| DCX | Rabbit | 1:500 | Abcam (ab18723) |
| Map2 | Mouse | 1:200 | Millipore (MAB3418) |
| NeuN | Mouse | 1:500 | Millipore (MAB377) |
| GFAP | Rabbit | 1:1000 | Millipore (AB5804) |
| Ki67 | Rabbit | 1:500 | ThermoFisher (PA5-19462 |
| Cleaved Caspase 3 | Rabbit | 1:500 | Cell signaling technologies (9661T) |
| cFOS | Rabbit | 1:500 | Cell signaling technologies (4384) |
| Anti-Rabbit IgG Alexa Fluor 488 | Donkey | 1:500 | Invitrogen (A-21026) |
| Anti-Mouse IgG Alexa Fluor 594 | Goat | 1:500 | Invitrogen (A-11032) |

**Supplementary Table 2: PCR primers used:**

| Target | Sequence | Efficiency | Reference |
|---|---|---|---|
| Human GAPDH | Fwd: TGCACCACCAACTGCTTAGC<br>Rvse: GGCATGGACTGTGGTCATGAG | 110% | Primer BLAST |
| Human HPRT | Fwd: AGCTTGCTGGTGAAAAGGAC<br>Rvse: TTATAGTCAAGGGCATATCC | 105.8% | Primer BLAST |
| Human RMST | Fwd: GCAGTGGGTGACTGATCGTA<br>Rvse: AGTCAACTCCGTGTCCCTTG | 97.7% | Primer BLAST |
| miR-125b | CTCCCTGAGACCCTAACTTGTG | 94.6% | IDTQuanta |
| miR-132 | CAGTCTACAGCCATGGTCGAAA | 109.3% | IDTQuanta |
| miR-135a | GCTATGGCTTTTTATTCCTATGTGA | 106.9% | IDTQuanta |
| miR-139 | CAGTGCACGTGTCTCCAGTAAAA | 108.5% | IDTQuanta |
| miR-146a | TGAGAACTGAATTCCATGGGTTA | 130.9% | IDTQuanta |
| miR-17 | CCACAAAGTGCTTACAGTGCAG | 108.7% | IDTQuanta |
| miR-181c | AACATTCAACCTGTCGGTGAGT | 107.4% | IDTQuanta |
| miR-19a | TGTGCAAATCTATGCAAAACTGA | 103.6% | IDTQuanta |
| miR-21 | GCTAGCTTATCAGACTGATGTTGAAA | 106.1% | IDTQuanta |
| miR-30a | CGATGTAAACATCCTCGACTGG | 104.6% | IDTQuanta |
| miR-9 | CGCTCTTTGGTTATCTAGCTGTATG | 97.2% | IDTQuanta |
| Pri-miR-135a1 | Fwd: CCTCGCTGTTCTCTATGGCTTT<br>Rvse: ACGGCTCCAATCCCTATATGA | 102.3% | Primer BLAST |
| Pri-miR-135a2 | Fwd: TCACTCTAGTGCTTTATGGCTT<br>Rvse: TGGCTTCCATCCCTACATGA | 101.4% | Primer BLAST |
| Universal primer | GCATAGACCTGAATGGCGGTA | | IDTQuanta |
